# Ultrahigh-Throughput Ambient MS: Direct Analysis at 22 Samples per Second by Infrared Matrix-Assisted Laser Desorption Electrospray Ionization Mass Spectrometry

**DOI:** 10.1101/2021.10.25.465730

**Authors:** Andrew J. Radosevich, Fan Pu, David Chang-Yen, James W. Sawicki, Nari N. Talaty, Nathaniel L. Elsen, Jon D. Williams, Jeffrey Y Pan

**Affiliations:** AbbVie Inc., 1 North Waukegan Rd., North Chicago, IL 60064

## Abstract

Infrared Matrix-Assisted Laser Desorption Electrospray Ionization (IR-MALDESI) mass spectrometry is an ambient-direct sampling method being developed for high-throughput, label-free, biochemical screening of large-scale compound libraries. Here, we report the development of an ultrahigh-throughput continuous motion IR-MALDESI sampling approach capable of acquiring data at rates up to 22.7 samples per second in a 384-well microtiter plate. At top speed, less than 1% analyte carryover is observed from well-to-well and signal intensity relative standard deviations (RSD) of 11.5% and 20.9% for 3 μM 1-hydroxymidazolam and 12 μM dextrorphan, respectively, are achieved. The ability to perform parallel kinetics studies on 384 samples with ~30s time resolution using an isocitrate dehydrogenase 1 (IDH1) enzyme assay is shown. Finally, we demonstrate the repeatability and throughput of our approach by measuring 115,200 samples from 300 microtiter plate reads consecutively over 5.54 hours with RSDs under 8.14% for each freshly introduced plate. Taken together, these results demonstrate the use of IR-MALDESI at sample acquisition rates that surpass other currently reported direct sampling mass spectrometry approaches used for high throughput compound screening.

**For Table of Contents Only:** 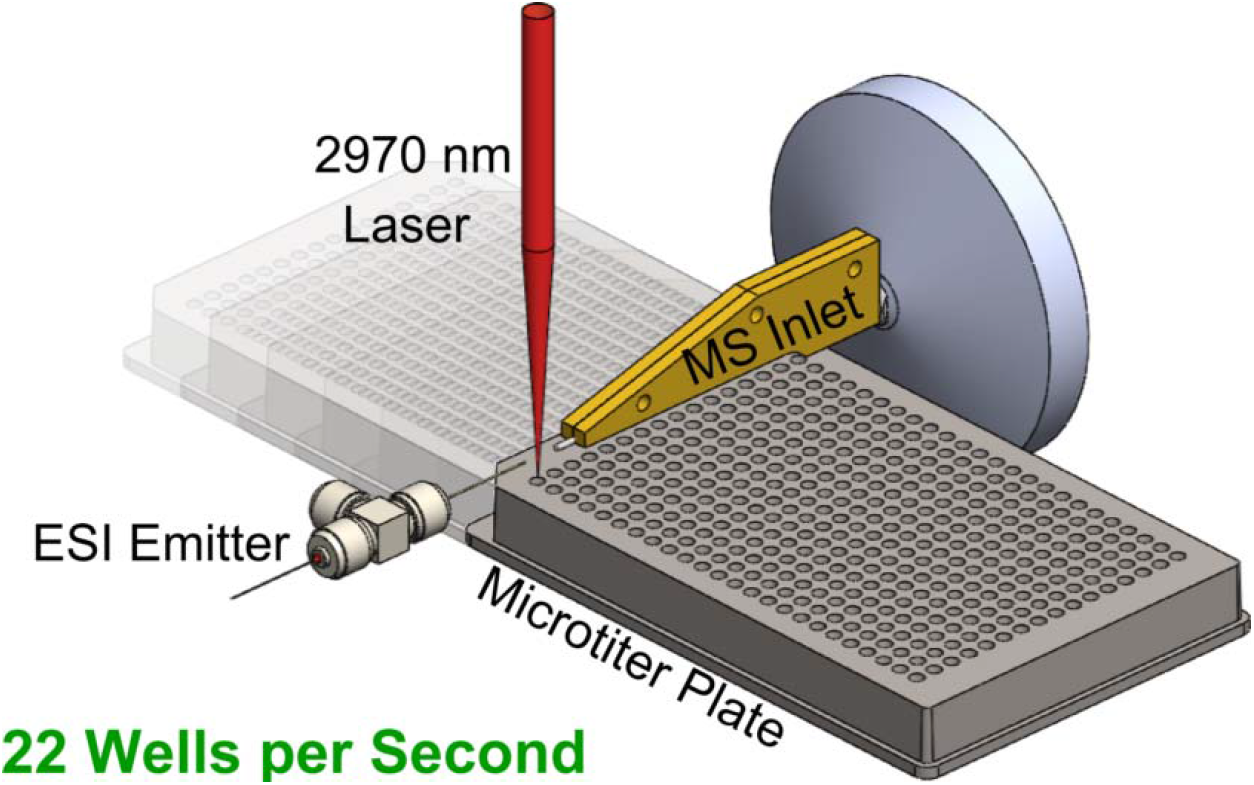

## Introduction

Adoption of mass spectrometry for ultrahigh throughput screening (uHTS) assays requires robust instrumentation capable of data collection at sample rates greater than 1 Hz since large pharmaceutical compound collections typically contain millions of compounds assayed in each biochemical screen. MALDI-MS can analyze multiple samples per second but requires addition of an organic matrix to form crystals, requiring an additional sample handling step that limits the ability to do large-scale real-time kinetics experiments efficiently^1^. Other suitable interfaces capable of sub-second per sample acquisition include acoustic mist (AMI-MS)^2, 3^, acoustic droplet ejection – open port interface (ADE-OPI)^4, 5^, and desorption electrospray ionization (DESI)^6^. To our knowledge, the fastest sampling rate of 6 Hz was achieved with ADE-OPI connected to a triple quadrupole MS system^7^.

Recently, we have developed IR-MALDESI as an alternative MS interface technology capable of high throughput direct analysis from any conventional microtiter plate format^8, 9^. Using a Q Exactive HF-X mass spectrometer, our group found a maximum potential spectra scan rate of 33 Hz at an MS resolving power (RP) of 7500 at m/z=200^9^. However, due to limiting factors such as the laser pulsing rate and energy stability, the speed and repeatability of the scan stage, and non-optimized control software, the practical scan speed achieved in a microtiter plate was much slower at 1.28 Hz or ~5 min for a full 384-well plate.

To overcome these limitations, we have made hardware and software improvements to maximize the sampling speed. In this work, we begin by describing updates to the mid-infrared ablation laser module and sample stage. We then detail the development of an ultrahigh-throughput continuous motion sampling approach capable of achieving a maximum of 22.7 Hz spectra scan rate which we refer to colloquially as Ludicrous Mode. Next, we use staggered plating of dextrorphan and 1-hydroxymidazolam to study the maximum scan rates and sample carryover at each MS RP. We then show the ability to run 384-well parallel kinetics studies with high temporal resolution (i.e., ~30 second intervals between measurements of the same well) using an isocitrate dehydrogenase 1 (IDH1) enzyme assay. We finish with a throughput demonstration by measuring 115,200 samples from 300 microtiter plates consecutively over 5.54 hours.

## Materials and Methods

### IR-MALDESI Instrumentation

Our IR-MALDESI system was previously described in detail^9^. In brief, a pulse of energy from a 2970 nm laser ejects a plume of droplets from a water matrix sample into an electrospray stream that ionizes the droplets as they move into the transfer tube of a Q Exactive HF-X (Thermo Fisher) mass spectrometer. A custom MS ion transfer tube with 3.14” length and a cartridge heater enable measurement from any standard 96, 384, or 1536 SBS format microtiter plate. The tip of the ion transfer tube was located 8 mm from the tip of the ESI capillary and 6 mm from the laser beam axis. The laser focal point was focused to the sample surface at a vertical distance of 5 mm from the center of the ion transfer tube axis. All plates are presented to the mass spectrometer in landscape format to minimize the length of the transfer tube. Scans follow a serpentine pattern that translates across the numbered columns of the plate before stepping to the next lettered row. A custom C# software coordinates device parameter setting, plate scanning, and syncing with the MS. Figure 1A shows an image of the IR-MALDESI source. Annotated figures of the full IR-MALDESI system can be found in Figure S1-S3.

**Figure 1.**
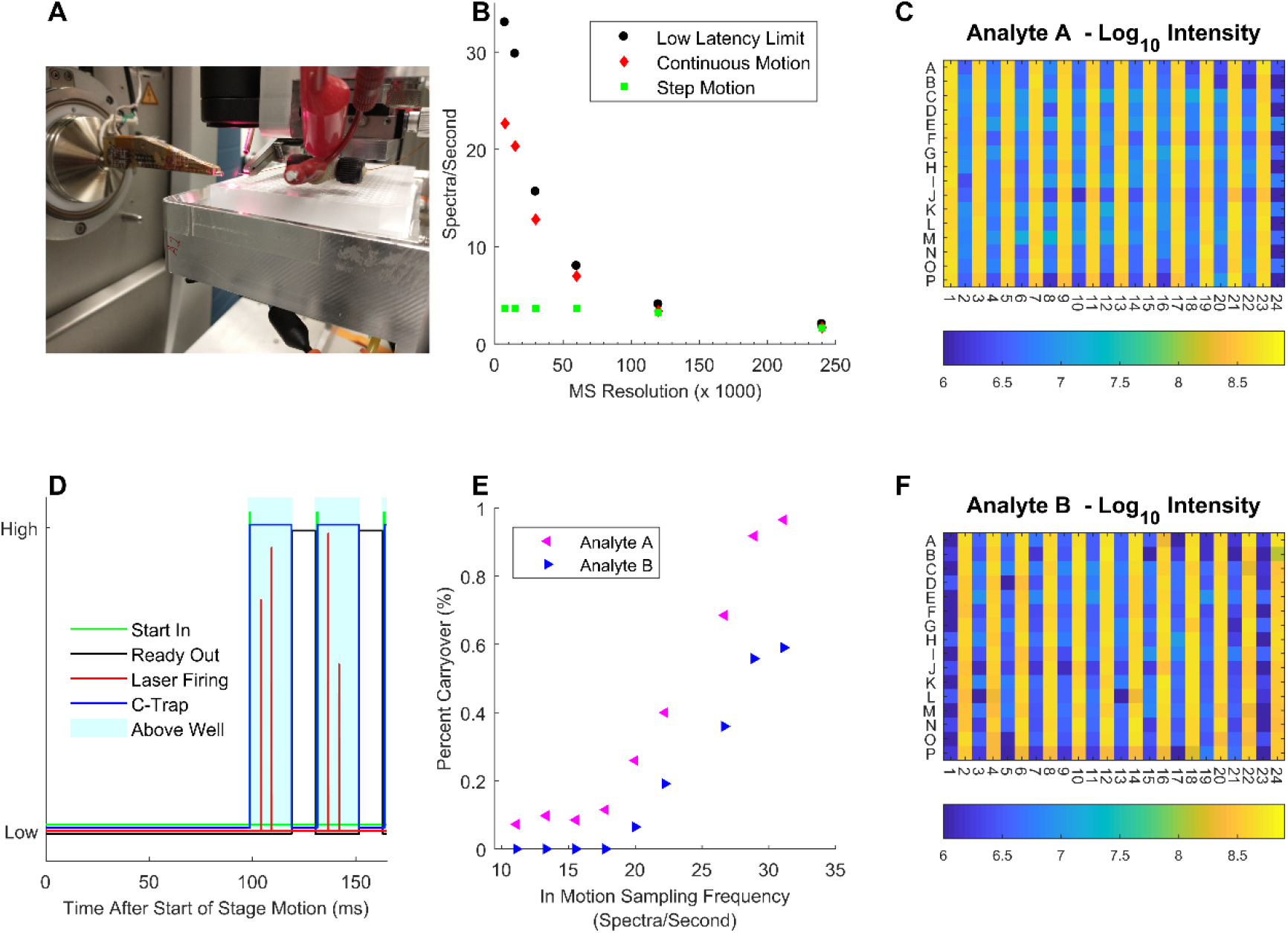
Demonstration of ultra-high speed acquisition in a continuous motion scan mode. A) Image of the IR-MALDESI source. B) Acquisition speed comparison between various data sampling scan modes. D) Communication signals for the first 2 spectra acquired at 7500 RP and 20 ms C-trap injection time. E) Percent carryover of analyte signal into subsequent well as a function of in motion sampling rate. C) and F) Plate maps with minimal sample carryover in continuous motion scan mode at 22.7 Hz for 12 μM dextrorphan and 3 μM 1-hydroxymidazolam, respectively.

To improve the speed and stability of the laser emission, we added active temperature control by modifying the top enclosure plate of the pulsed mid-IR laser (JGMA Inc.) detailed previously^10^. The temperature directly above the active portion of the laser was maintained at 25 °C by a fan-cooled Peltier chiller module (Adafruit, product ID: 1335) and thermoelectric cooler controller (Thorlabs, TED4015). With this custom configuration, we can reach pulse rates of 100 Hz or 5 times faster than the 20 Hz provided by the leading commercial option of the Opolette OPO laser by Opotek. For each MS spectrum, the laser fires two bursts of light separated by 5 ms, with each burst composed of 2 pulses separated by ~60 microseconds for a total energy of ~2 mJ (see Figure S4-S6).

To improve speed, repeatability, and accuracy of our stage movement, we replaced the primary axis (i.e., the horizontal axis parallel to the front panel of the MS) stepper motor drive with a 100W servo motor with a high-resolution encoder (J0100-303-3-000, Applied Motion Products). The position of the servo motor is polled via RS-232 communication at an interval of ~16 ms with intermediate positions extrapolated in software at better than 100 μs resolution.

### Continuous motion scanning

A common issue limiting the speed of scanning stage-based instruments is an inability to rapidly move the stage between static locations to sample data. To overcome this issue, we developed a continuous motion scan mode in which all data is collected while the scan stage is in motion over the top of sample wells (Videos S1 and S2). To avoid the acceleration phase of the servo motor motion, each X-axis scan begins 4.5 mm (i.e., a single 384-well separation) before the first well and finishes 4.5 mm after the final well in each row.

Figure 1D shows the timing of communication between the MS, X-axis scan stage and laser firing for the first several spectra acquired at 7500 RP. During the first 90-100 ms, the scan stage accelerates to its maximum speed while the control computer continuously polls its position. When a well moves under the MS sampling point (as depicted in the shaded region), a start-in contact closure is sent to the MS to request an acquisition. Within 300 μs, the split lens opens the C-trap for ion accumulation over the injection time specified in the Xcalibur method. Two bursts of mid-IR energy are delivered to the sample 5 ms after the C-trap opens, thereby introducing sample analytes into the electrospray for eventual capture in the C-trap. As the ions are being read in the Orbitrap, the split lens closes ion introduction into the C-trap and the MS firmware blocks sampling until the ready-out contact closure is opened, at which point a new acquisition can be triggered. For different RP, the communication sequence looks similar except that the length of time the ready-out signal remains high increases for higher RP values (Figure S5).

The linear speed of the servo motor is set such that the frequency at which wells fall under the MS sampling point is approximately the maximum MS sampling frequency for a given RP. To allow for slight variability in the length of timing events, the selected speed will always be lower than the theoretical max. Column 2 of Table S1 shows the actual linear scan speeds used in our experiments for acquisitions at different RP.

### Mass Spectrometer Settings

Experiments were carried out with a 3.7 kV spray voltage in positive ion mode. The ion accumulation time was 20 ms with a scan range of positive *m/z* 200-400 (for dextrorphan and 1-hydroxymidazolam) or negative *m/z* 100-210 (for IDH1 assay). The ion transfer tube temperature was maintained at 400°C. The ESI solvent of 80:20 methanol/water with 0.1% formic acid was sprayed at a constant flow rate of 2 μL/min throughout an experiment.

### IDH1 Kinetics Assay

The assay buffer contained 5 mM Tris pH 7.5, 1 mM MgCl_2_, 1 mM DL-Dithiothreitol, and 0.01 wt. % bovine serum albumin. Substrate concentrations were 100 μM isocitrate and 40 μM for NADP^+^. A titration of IDH1 enzyme was plated across subsequent sets of 6 columns in a 384-well microtiter plate as shown in Figure 2A. An equal volume of substrate and enzyme solution was mixed to a volume of 20μL with final concentrations of 0 nM, 1.56 nM, 3.125 nM, 6.25 nM, 12.5 nM and 25 nM. The reaction was carried out at room temperature. The first MS measurement in the kinetics scan was taken 15 seconds after addition of substrate solution to all wells. A full 384-well plate worth of data was collected in ~30 seconds with a RP of 30 000 (FWHM at m/z = 200). 10 full scans were repeated back-to-back with a 3 seconds delay between scans to allow the stage to return to its starting point.

**Figure 2.**
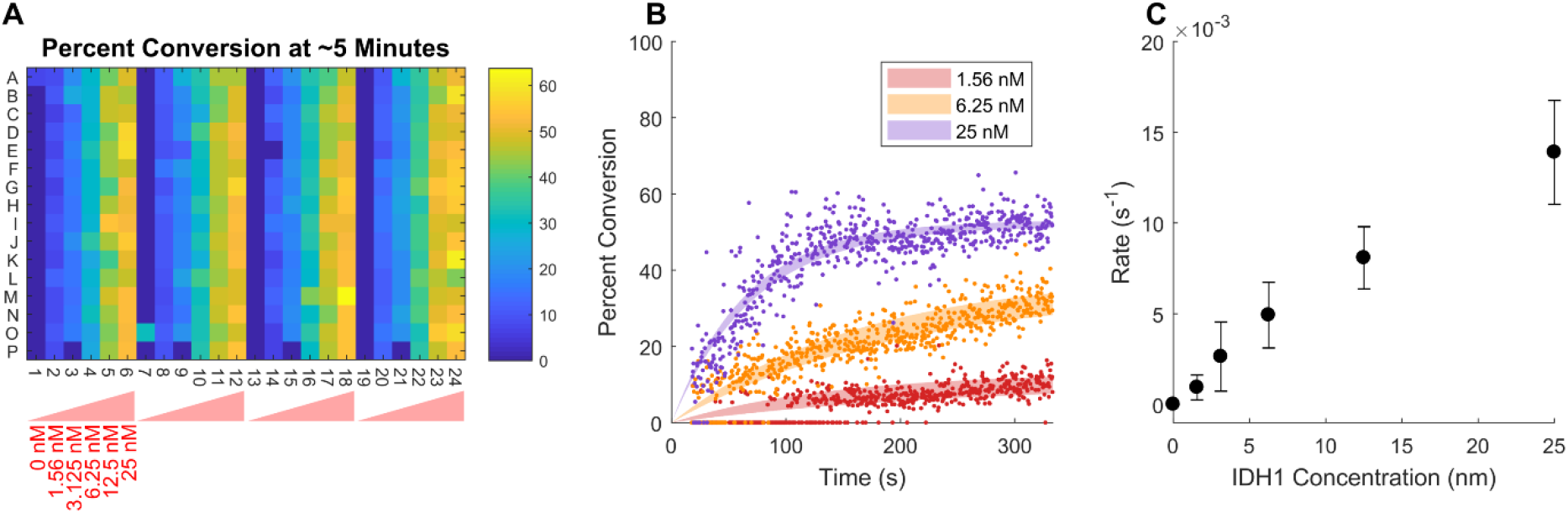
Measuring enzyme kinetics on 384 samples in parallel. Same well time resolution is ~30 seconds at 30000 RP. A) Plate map of IDH1 assay percent conversion at the 5-minute time point. B) Percent conversion as a function of measurement time for all wells and time points. Shaded region shows the 95% confidence interval on the curve fit. C) Rate of percent conversion as a function of IDH1 enzyme concentration. Error bars show standard deviation across wells.

### Ultra-high Throughput Measurement Demonstration

Three 384-well plates mimicking a reaction quenched timepoint of the IDH1 assay were prepared with alternating columns of 50 μM isocitrate + 50 μM α-ketoglutarate in assay buffer at a well volume of 20μL. 300 full plate scans were run consecutively on the same plate except for brief delays to switch to fresh plates at the 101^st^ and 201^st^ scans. A single plate scan consisted of ~30 seconds of data collection (30 000 RP; FWHM at m/z=200) followed by a 30 second delay to simulate time for robotic plate switching.

### Data Processing and Analysis

Data processing was done using a previously described in-house data browser^9^. Model fitting and other plots were made with MATLAB version R2017a. Percent carryover was measured as:

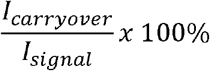

where *I*_*signal*_ is the mean intensity of wells containing the analyte of interest and *I*_*carryover*_ is the mean intensity of wells not containing the analyte of interest. Percent conversion for the IDH1 assay was calculated as:

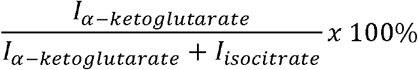

where *I* is the ion abundance of α-ketoglutarate (m/z 145.013) and isocitrate (m/z 191.018) summed within a 10 ppm window. Enzyme kinetics for the conversion of isocitrate to α-ketoglutarate were modeled as:

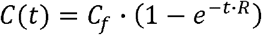

where C(t) is the percent conversion as a function of time, *C*_*f*_ is the final reaction percent conversion and R is the rate of reaction.

## Results and Discussion

### Continuous motion scan speeds and data quality

Figure 1B shows the acquisition speeds obtained under continuous and step motion scan modes against the maximum sampling rate possible using a low latency handshake. At 120 000 and 240 000 RP, the rate limiting component is the sampling rate of the MS. As a result, both continuous and step motion scan modes closely approach the low latency handshake limit. As RP decreases, the acceleration/deceleration profile of the scan stage becomes limiting under a step motion scan mode, resulting in a plateaued 3.66 sample per second scan rate for RP under 60 000 (column 5 of Table S1). In this regime, the continuous motion scan mode continues to exhibit increased scan rates with a maximum of 22.7 samples/second at 7500 RP (column 4 of Table S1).

The scan rates shown in Figure 1B are the average over the full 384-well plate (i.e., simply 384 divided by the total scan time). As RP decreases, this falls further away from the low latency handshake limit due to the increasing relative contribution of inefficiencies such as the stage acceleration phase, slight variability in acquisition syncing, and control software overhead. Neglecting the start and end of motion acceleration phases, sampling frequency reaches 31.11 Hz while the stage is at a constant velocity. The 3^rd^ column of Table S1 summarizes the sampling frequencies only considering the time while the stage is in motion.

To study the amount of sample carryover from well to well, we plated alternating columns with 12 μM dextrorphan and 3 μM 1-hydroxymidazolam. At the fastest sample rate at 7500 RP, less than 1% carryover is seen in the signal intensity plate maps for both dextrorphan (Figure 1C) and 1-hydroxymidazolam (Figure 1F). Furthermore, the signal intensity RSD across all wells was excellent for both dextrorphan (20.9%) and 1-hydroxymidazolam (11.5%). Although we use these small molecules as demonstration, minimal carryover and good quality spectra were also obtained for a much larger ~12 kDa cytochrome C molecule (Figure S 7).

To determine the effect of speed on carryover, we held the RP constant at 7500 and varied the sampling frequency by changing the stage linear velocity. For sampling frequencies higher than ~18 Hz, the percent carryover increases from under 0.1% up to ~1% for both analytes (Figure 1E). We attribute this increase in carryover to analyte from the previous ablation event remaining suspended in the sample plume. This will tend to occur when the sampling time is shorter than the plume lifetime of ~50 ms, corresponding to 20 Hz sampling frequency^11^. A similar trend is seen when varying sampling frequency by running the MS at its maximum collection rate at different RP (Figure S 8).

### Using ultrahigh scan speeds to perform rapid rate kinetics assays in parallel

One application enabled by ultrahigh scan rates is the ability to measure kinetics on the order of 10’s of seconds from a large array of samples in parallel. Figure 2 provides a demonstration of this capability using an IDH1 enzyme reaction. Figure 2A shows a plate map of the conversion of isocitrate to α-ketoglutarate for samples with varying concentration of IDH1 enzyme for the time point starting around 5 minutes. Replicates across the microtiter plate show qualitative agreement except for artifacts in the “P” row due to pipetting error. Figure 2B shows the change in percent conversion for each time point in the data set with the shaded regions indicating the 95% confidence interval of an exponential plateau curve fit. Finally, Figure 2C shows the mean and standard deviation of the rate term of the exponential plateau curve fit for each well.

### Ultrahigh-throughput measurement of more than 100,000 samples in under 6 hours

To demonstrate the robustness of IR-MALDESI under a continuous motion scan mode, we ran 300 consecutive 384-well plate measurements (115,200 total data points at 30 000 RP) over the course of 5.54 hours. Fresh plates were loaded onto the instrument at the 1^st^, 101^st^, and 201^st^ scans due to sample evaporation. Figure 3 shows qualitative agreement and minimal sample carryover between plate maps of IDH1 percent conversion for each scan where a fresh plate was loaded. For scans 1, 101, and 201, the mean percent conversion across sample wells was 57.0%, 59.6%, and 61.5% with RSDs of 6.09%, 7.92%, and 8.14%, respectively. The mean percent carryover across all blank wells for all scans was 0.17%.

**Figure 3.**
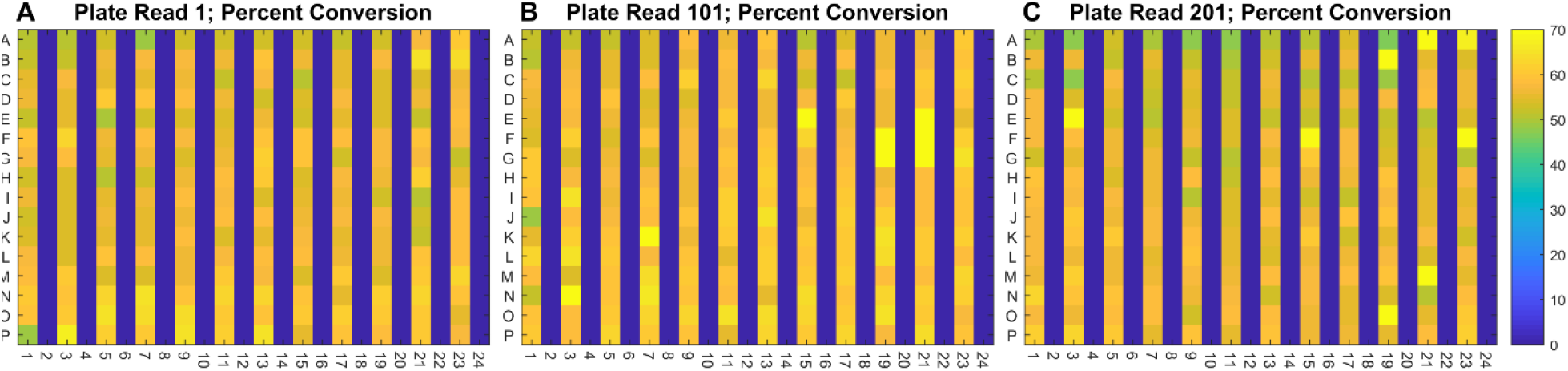
Demonstration of signal stability and minimal carryover across hundreds of scans. A), B) and C) show plate maps of the IDH1 percent conversion metric at timepoints 0 (Scan 1), 1.83 hours (Scan 101), and 3.87 hours (Scan 201), respectively.

Plots for the percent conversion, RSD, and percent carryover for each scan are provided in Figure S9. Percent conversion (minimum=56.3% and maximum=67.8%) and RSD (minimum=5.58% and maximum 16.22%) for the same plate increase over time but return to baseline when a fresh plate is introduced. We therefore attribute these changes to sample evaporation while the MS measurement remains robust. Mean percent carryover for each plate (minimum=0% and maximum 11.72%) is driven by a handful of wells, with 96.5% of all blank wells having isocitrate and α-ketoglutarate signal intensities of exactly 0. We attribute the rare high outlier carryover values to the random generation of large ablation droplets that temporarily contaminate the transfer tube or electrospray capillary (Figure S10).

## Conclusions

This report described the application of a continuous motion scan approach to acquiring ultra-high speed IR-MALDESI measurements on a Thermo Q Exactive HF-X MS. Importantly, this approach can be performed in conventional microtiter plates and is amenable to application on other types of MS. A maximum in-motion sampling frequency of 31.1 Hz at 7500 RP was reduced to 22.7 Hz due to the practicalities of measuring samples in a 384-well microtiter plate. As the RP increase, ions spend a longer time in the Orbitrap analyzer resulting in a decreased sampling frequency overall. When the RP reaches 120 000 or above, the MS acquisition is sufficiently longer than the stage single step settling time that there is no benefit to performing a continuous motion scan at these resolutions.

The benefit of such a high sampling frequency is optimized when as many tests as possible can be run for any sample plate loading event. This argues for the development of improved sample handling automation, use of high- or custom-density well plates, taking multiple replicates of the same plates for signal averaging, or the application of kinetic studies. Here, we demonstrated the ability to measure the kinetics of 384 samples in parallel with time resolution of ~30 seconds using an IDH1 enzyme assay. Furthermore, we demonstrated the repeatability of our approach by measuring 300 microtiter plates consecutively over the course of 5.54 hours. Importantly, despite introducing a high number of samples into our MS in such a short period of time, we saw no deterioration of performance or accumulation of carryover with increasing scan number. Future studies will assess how many samples can be analyzed successfully without cleaning the inlet and/or ion funnel.

The sampling frequencies presented in this work compare favorably against those of other direct sampling MS technologies such as AMI-MS at 3 Hz^3^, DESI at 3.3 Hz^6^, and ADE-OPI at 6 Hz^7^. Moreover, with total scan times under 20 seconds for a 384-well plate, IR-MALDESI reaches speeds rivalling those of a variety of optical technologies while retaining the advantage of true label-free screening. As a result, we envision the use of IR-MALDESI as an important tool for future high-throughput biochemical screens.

## Supporting information

Continuous Motion Scan Video

Software Interface Video

## Supporting Information

Additional experimental details including photographs and videos of experimental setup

## Acknowledgements

The authors acknowledge Måns Ekelöf (EMBL), Ken Garrard (NCSU), M. Caleb Bagley (NCSU), David C. Muddiman (NCSU), Jeffrey G. Manni Sr (JGM Associates, Inc.) and Kyle Fort (Thermo Fisher) for their critical insights. Funding was provided to David C. Muddiman (NCSU) by AbbVie. There is no funding to disclose for Måns Ekelöf (EMBL), Ken Garrard (NCSU), M. Caleb Bagley (NCSU), Jeffrey G. Manni Sr (JGM Associates, Inc.), and Kyle Fort (Thermo Fisher). The work was enabled by the AbbVie Postdoc Program. All authors are employees of AbbVie. The design, study conduct, and financial support for this research were provided by AbbVie. AbbVie participated in the interpretation of data, review, and approval of the publication.

