## Supporting Information for "Ultrahigh-Throughput Ambient MS: Direct Analysis at 22 Samples per Second by Infrared Matrix-Assisted Laser Desorption Electrospray Ionization Mass Spectrometry"

### IR-MALDESI Instrument Models and Description

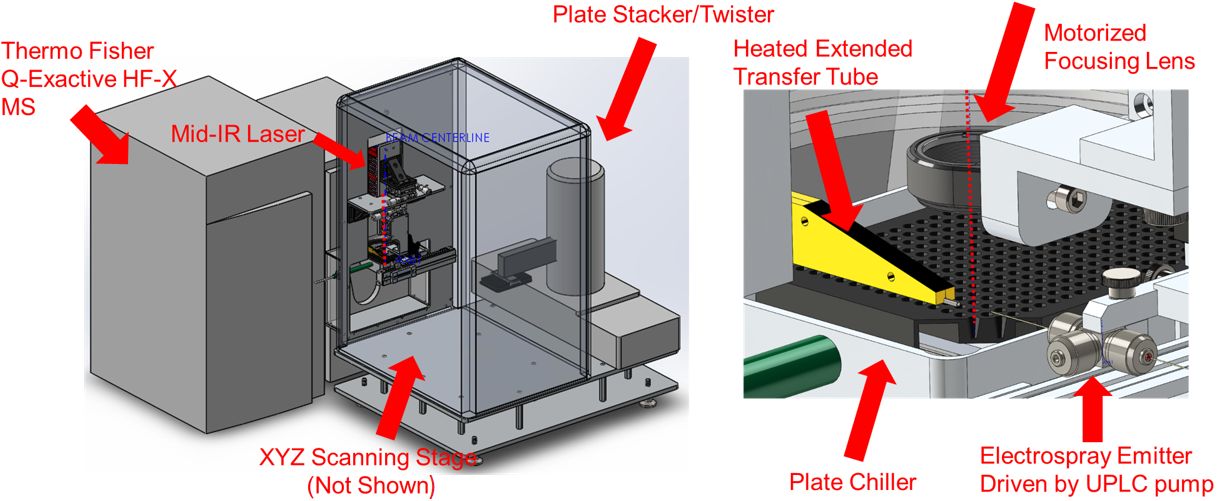

Figure S 1 3D solid model of the IR-MALDESI instrument and Q Exactive HF-X MS

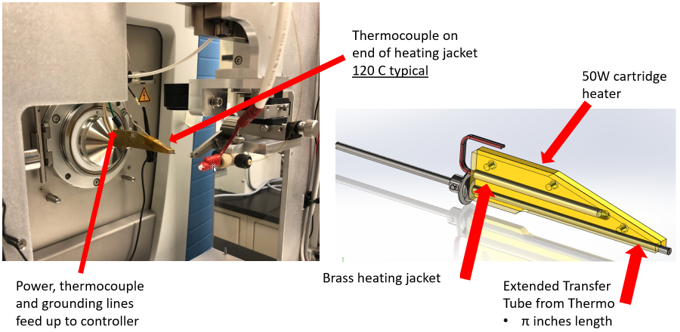

Figure S 2 Detail of the extended, heated ion transfer tube

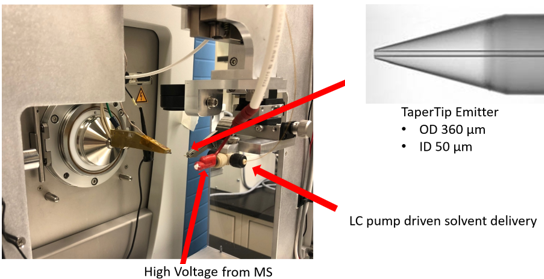

Figure S 3 Detail of the electrospray ionization source

### Coordination of acquisition signals between MS and custom instrument control software

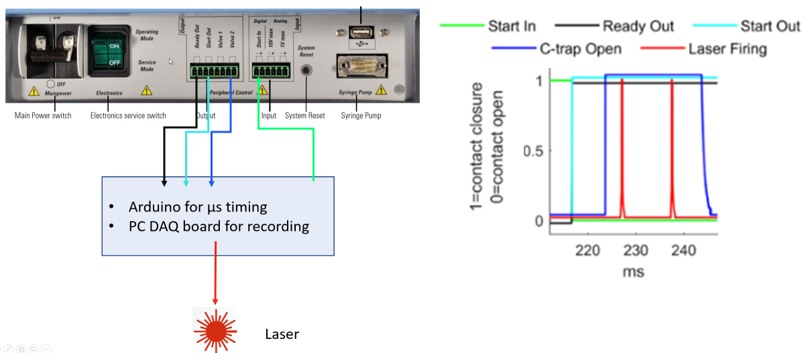

Figure S 4 Diagram showing communication signals coming into and out of the Q Exactive HF-X MS Instrument Panel. The Ready Out signal is a contact closure output that opens when the MS is ready to acquire a new spectrum. The Start In signal is a contact closure input that is closed by our custom Arduino program to trigger a new spectrum acquisition. The Start Out signal is a contact closure output that opens when a new spectrum acquisition has started. The C-trap Open signal is a contact closure output that opens when the C-trap is accumulating ions. The red laser firing trace is an analog signal from a photodiode embedded in the mid-IR ablation laser. Custom C# control software interacts with an Arduino microprocessor to coordinate sample scanning and MS acquisition.

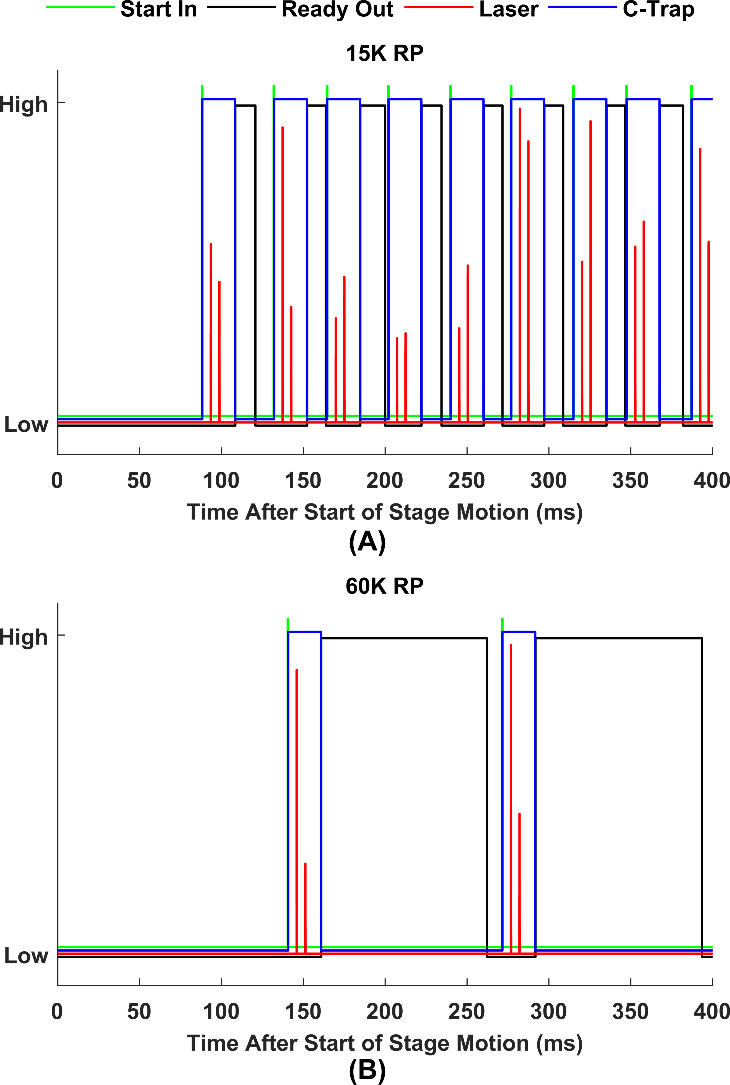

Figure S 5 Communication signals for continuous motion scan at 15 000 RP (A) and 60 000 RP (B)

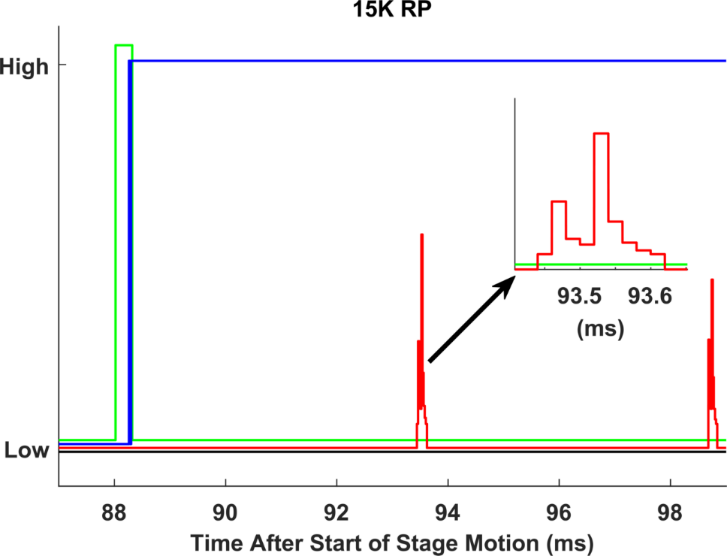

Figure S 6 Zoomed in communication signal showing two laser pulses for each of the two laser bursts.

### Summary of acquisition speed values and video demonstration

Table S 1 Scan speeds in a 384-well plate. In motion sampling frequency gives the scan rate between wells neglecting start/end of motion delays. Effective continuous motion sampling frequency is the average scan rate across a full 384-well plate. Step motion sampling frequency is the scan rate when the stage stops at each well position before proceeding to the next well.

| Resolving Power | Continuous Motion Scan Stage Speed (mm/s) | In Motion Sampling Frequency (Hz) | Effective Continuous Motion Sampling Frequency (Hz) | Step Motion Sampling Frequency (Hz) |
| --- | --- | --- | --- | --- |
| 7500 | 140 | 31.11 | 22.67 | 3.66 |
| 15000 | 120 | 26.67 | 20.34 | 3.66 |
| 30000 | 68 | 15.11 | 12.81 | 3.66 |
| 60000 | 35 | 7.78 | 6.98 | 3.66 |
| 120000 | 17.5 | 3.89 | 3.38 | 3.25 |
| 240000 | 9 | 2 | 1.7 | 1.6 |

*Video S 1 Video of instrument scan stage motion while acquiring at 22.7 Hz using a continuous scan motion at 7500 RP. The first 4.5 seconds are real-time while the remaining seconds are in slow motion to demonstrate the laser pulse synchronization with each well location. The continuous red laser enables us to visualize the electrospray while the intermittent yellow flashes are the mid-IR desorption laser hitting the middle of each well.*

*Video S 2 Video showing real-time scan progress in our custom software (i.e., red to green square color change) alongside the spectra shown in Xcalibur for acquisition at 22.7 Hz using a continuous scan motion at 7500 RP. The sample was alternating columns of 0.13 mg/mL cytochrome C in water. The processed data from this experiment are shown in Figure S 7.*

### Demonstrations of carryover and signal robustness under a continuous motion scan

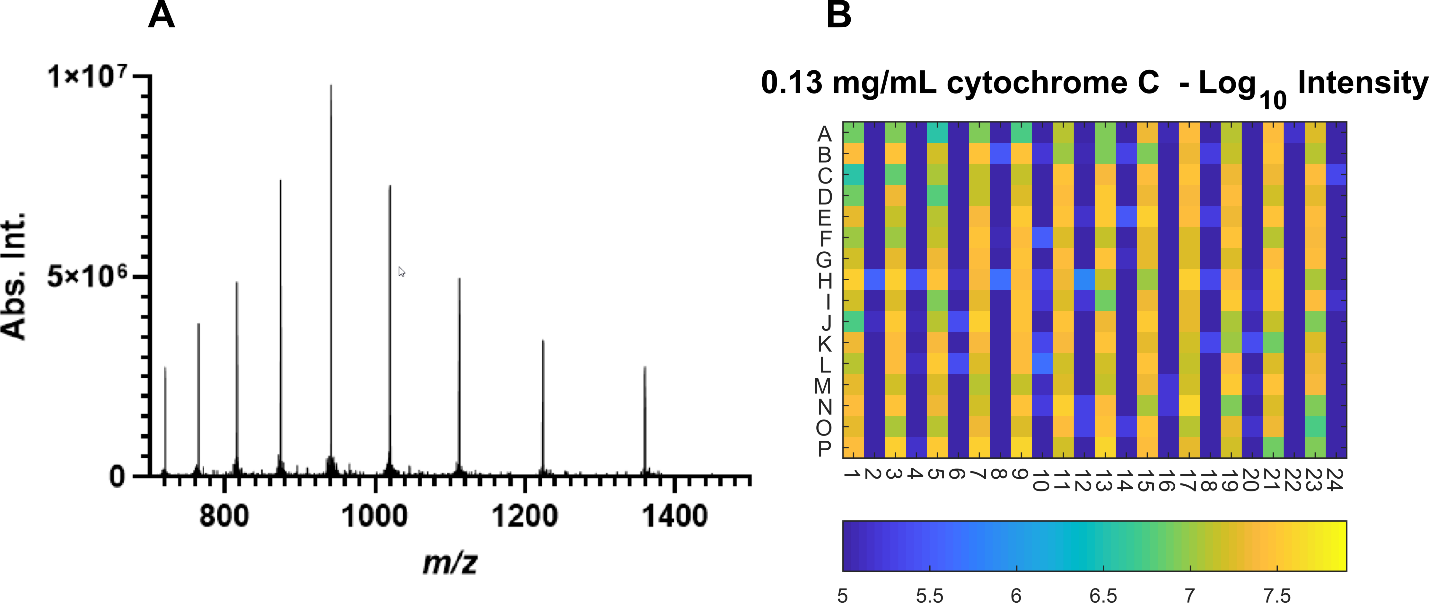

Figure S 7 Demonstration of continuous motion scan mode at 22.7 Hz for high molecular weight (~12 kDa cytochrome C at 0.13 mg/mL in water) analyte. A) example spectrum for cytochrome C and B) heat map showing minimal carryover to blank wells.

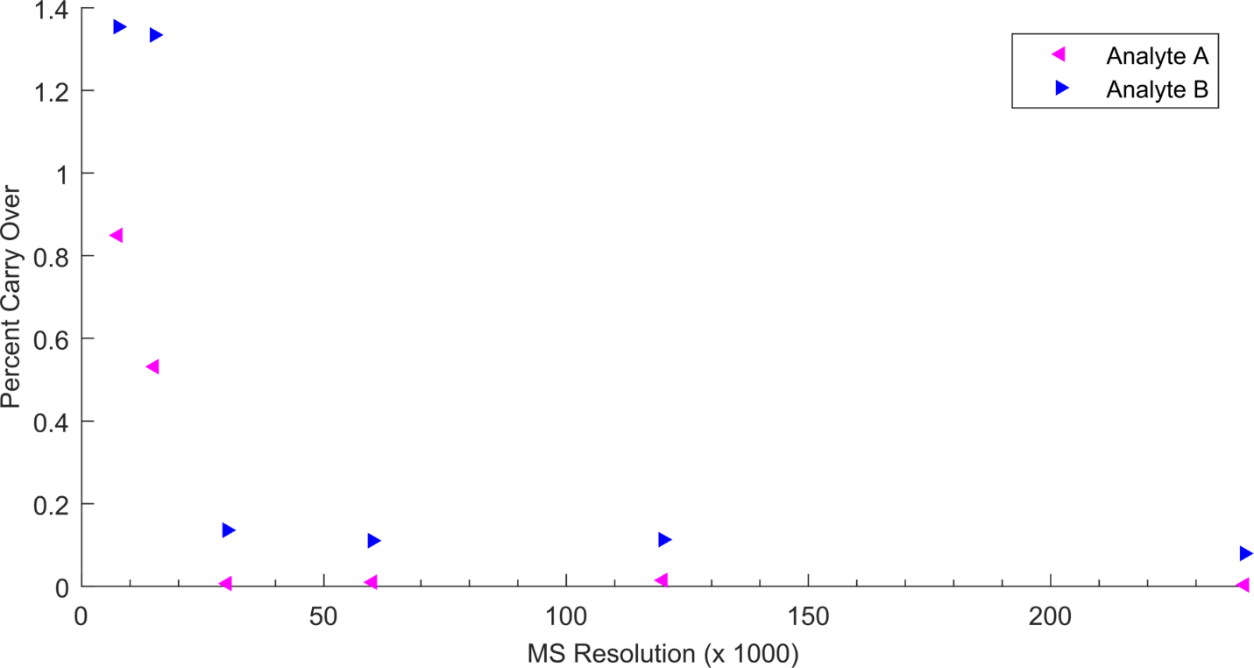

Figure S 8 Percent carryover between wells at different RP for 12 µM dextrorphan (Analyte A) and 3 µM 1-hydroxymidazolam (Analyte B)

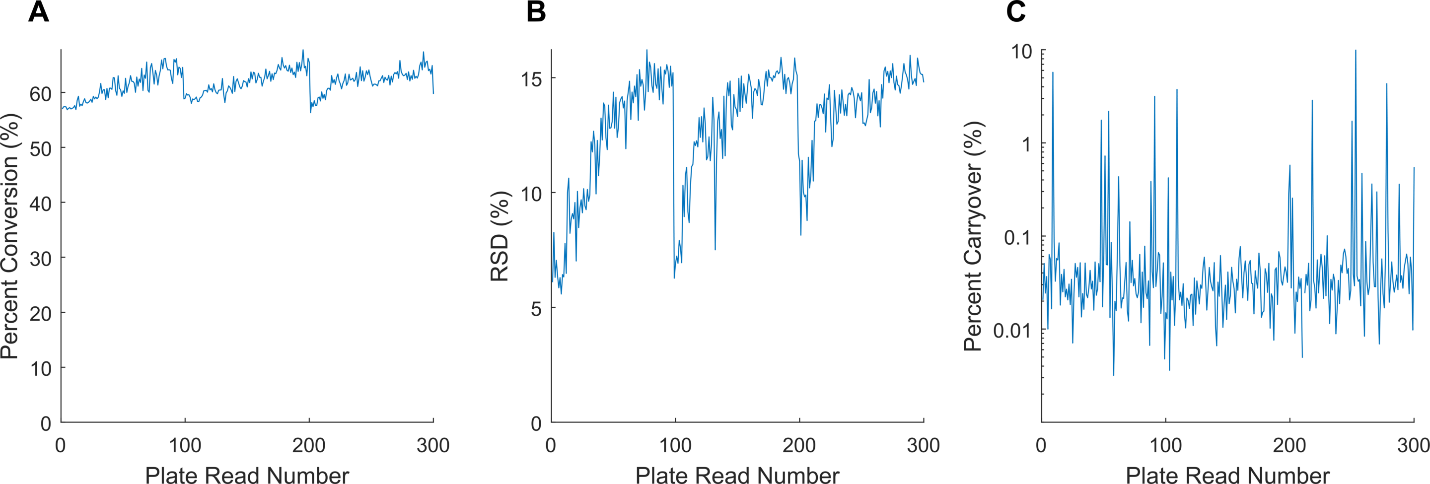

Figure S 9 Summary metrics from 300 consecutive plate scans under a continuous scan motion at 30 000 RP. Each scan consisted of 30 seconds of data acquisition followed by 30 seconds of delay to simulate robotic plate switching. Total scan time was 5.54 hours. Fresh sample plates were introduced at scan numbers 1, 101, and 201. A) Average percent conversion (α-ketoglutarate/(isocitrate+α-ketoglutarate)*100) over all sample wells for each scan, B) Average RSD of the percent conversion values over all sample wells for each scan, C) Total percent carryover of α-ketoglutarate + isocitrate signal intensites into blank wells for each scan.

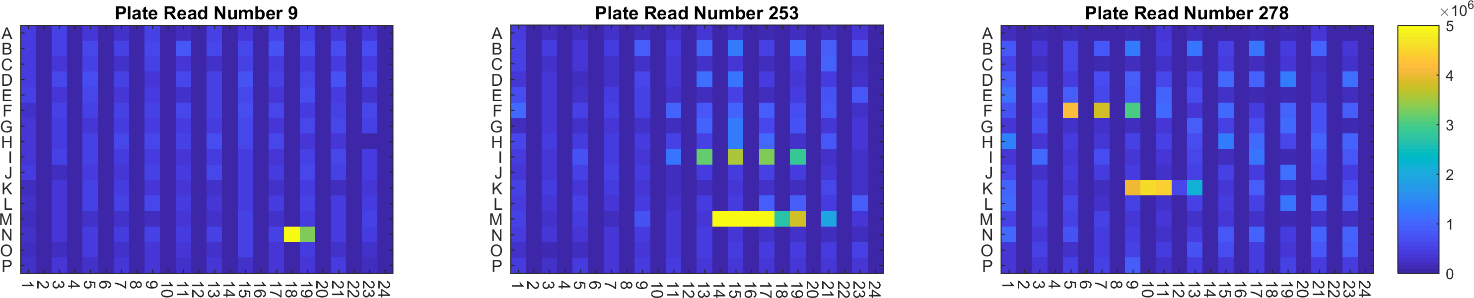

Figure S 10 Selected α-ketoglutarate intensity heat map examples of high carryover value scans. Although most blank wells have exactly 0 signal intensity, a handful of aberrantly high intensity wells drive the overall carryover values. We attribute these artifacts to random large ablation droplets that temporarily contaminate the electrospray capillary or ion transfer tube. In the worst cases as demonstrated above, the aberrant signal persists over multiple wells.
